# Modelling drug responses and evolutionary dynamics using triple negative breast cancer patient-derived xenografts

**DOI:** 10.1101/2023.01.10.523259

**Authors:** Abigail Shea, Yaniv Eyal-Lubling, Daniel Guerrero-Romero, Raquel Manzano Garcia, Wendy Greenwood, Martin O’Reilly, Dimitra Georgopoulou, Maurizio Callari, Giulia Lerda, Sophia Wix, Agnese Giovannetti, Riccardo Masina, Elham Esmaeilishirazifard, Alistair G. Martin, Ai Nagano, Lisa Young, Steven Kupczak, Yi Cheng, Helen Bardwell, Elena Provenzano, Justine Kane, Jonny Lay, Louise Grybowicz, Karen McAdam, Carlos Caldas, Jean Abraham, Oscar M Rueda, Alejandra Bruna

**Affiliations:** Cancer Research UK Cambridge Institute and Department of Oncology, Li Ka Shing Centre, University of Cambridge, Cambridge, UK; Centre for Paediatric Oncology Experimental Medicine, Centre for Cancer Evolution: Molecular Pathology Division, Institute of Cancer Research, Sutton, UK; Cambridge University NHS Foundation Trust, Cambridge, UK; Precision Breast Cancer Institute, Department of Oncology, University of Cambridge, Cambridge, UK; Fondazione Michelangelo, Milan; Clinical Genomics Unit, Fondazione IRCCS Casa Sollievo della Sofferenza, San Giovanni Rotondo (FG), Italy; MRC-Biostatistics Unit, University of Cambridge, Cambridge, United Kingdom; Addenbrookes Hospital, Department of Histopathology; NIHR Cambridge Biomedical Research Centre

## Abstract

Triple negative breast cancers (TNBC) exhibit inter- and intra-tumour heterogeneity, which is reflected in diverse drug responses and interplays with tumour evolution. Here, we use TNBC patient-derived tumour xenografts (PDTX) as a platform for co-clinical trials to test their predictive value and explore the molecular features of drug response and resistance. Patients and their matched PDTX exhibited mirrored drug responses to neoadjuvant therapy in a clinical trial. In parallel, additional clinically-relevant treatments were tested in PDTXs *in vivo* to identify alternative effective therapies for each PDTX model. This framework establishes the foundation for anticipatory personalised therapies for those patients with resistant or relapsed tumours. The PDTXs were further explored to model PDTX- and treatment-specific behaviours. The dynamics of drug response were characterised at single-cell resolution revealing a novel mechanism of response to olaparib. Upon olaparib treatment PDTXs showed phenotypic plasticity, including transient activation of the immediate-early response and irreversible sequential phenotypic switches: from epithelial to epithelial-mesenchymal-hybrid states, and then to mesenchymal states. This molecular mechanism was exploited *ex vivo* by combining olaparib and salinomycin (an inhibitor of mesenchymal-transduced cells) to reveal synergistic effects. In summary, TNBC PDTXs have the potential to help design individualised treatment strategies derived from model-specific evolutionary insights.

## INTRODUCTION

Breast cancers exhibit extensive inter- and intra-tumour heterogeneity at both the genomic and phenotypic levels. This is reflected in the diversity of interpatient drug responses, clinical prognoses and patterns of relapse (1–9). Patient-derived tumour xenografts (PDTXs) have been shown to retain both the inter-patient diversity and intra-tumour heterogeneity, and offer an unprecedented opportunity to study human breast cancer at high resolution (10–18). We have previously generated and molecularly characterised a living biobank of breast cancer PDTXs and demonstrated that these recapitulate the main histological and genomic features of the originating tumour, including intra-tumour heterogeneity (19). In addition, we pioneered the use of dissociated PDTX cells (PDTCs) for high-throughput drug screening, with most drug responses validated *in vivo*. Recently, we developed a panel of antibodies for mass-cytometry tailored to breast cancer PDTXs, allowing highly resolved multiparametric single cell phenotyping and its impact on drug responses (20).

The extent to which PDTXs mirror therapy responses of the matched donor patient has been insufficiently studied. This is further complicated when patients receive multiple lines of therapy prior and subsequent to the development of the PDTX model. This has limited the use of PDTXs as anticipatory tools and a systematic assessment of models from treatment-naïve patients in a controlled setting is required to simulate clinical practice.

Triple negative breast cancers (TNBC) generally have a poor prognosis, lack of actionable targets (21, 22) and are highly heterogeneous, impeding the development of improved therapeutic strategies (23–27). Advances in our understanding of the DNA damage response (DDR) offer new therapeutic avenues. PARP inhibitors (PARPi), e.g. olaparib, have a synthetic lethal effect in tumours with *BRCA1/2* alterations (28, 29) and have demonstrated clinical utility for patients who harbour these alterations (30–32). However, as with all targeted therapies, drug resistance and disease relapse remain obstacles. While several PARPi resistance mechanisms have been described (33), preclinical resistance studies typically have limited relevance to the clinical setting.

Here we have developed and optimised a PDTX platform alongside an ongoing neoadjuvant clinical trial for patients with TNBC and/or *BRCA1/2* alterations. Using mirrored treatment schedules of chemotherapy with or without olaparib, we show that PDTXs in a co-clinical trial are excellent laboratory model systems to reflect clinical drug responses. From this strong foundation, we expanded the potential of the PDTX platform by using it to test alternative treatment strategies to that which the given patient received. This enabled us to empirically and functionally identify best drug response data, and map at high resolution molecular landscape dynamics to retrieve evolutionary information. A deep-dive analysis into one PDTX model at single-cell resolution (using both single cell mass cytometry and single-cell RNA-sequencing) revealed a permanent phenotypic (non-genetic) switch to a mesenchymal cell state upon treatment with olaparib, which could be synergistically targeted with salinomycin, offering new avenues for therapeutic intervention in TNBC.

Our study has profound implications for the future use of PDTXs as an avatar platform and its use for anticipatory and predictive drug testing in the clinical setting, and a means to accelerate drug development with improved clinical predictive power.

## RESULTS

### A cohort of triple-negative breast cancer (TNBC) patient-derived tumour xenografts (PDTXs)

Patients with TNBC and/or germline alterations in *BRCA1/2* were enrolled on a neoadjuvant clinical trial, PARTNER (ClinicalTrials.gov Identifier: NCT03150576), which tested the efficacy of the PARPi olaparib in combination with two chemotherapy agents (carboplatin and paclitaxel). Tumour samples at diagnosis (therapy-naïve) from consenting patients were collected fresh and immediately transported to the animal facility for subcutaneous engraftment into NSG mice as previously described (19) (Figure 1A). Sixty-seven independent samples were implanted which resulted in 22 engrafted (grown as PDTX) and 14 established models (successfully passaged). This 33% engraftment rate is in line with that previously published by our group (19) and others (34). Tumour samples from the 14 that were successfully established as PDTXs (PDTX cohort) reflected the clinical characteristics of all 67 implanted samples (clinical trial cohort) (Figure 1B).

**Figure 1:**
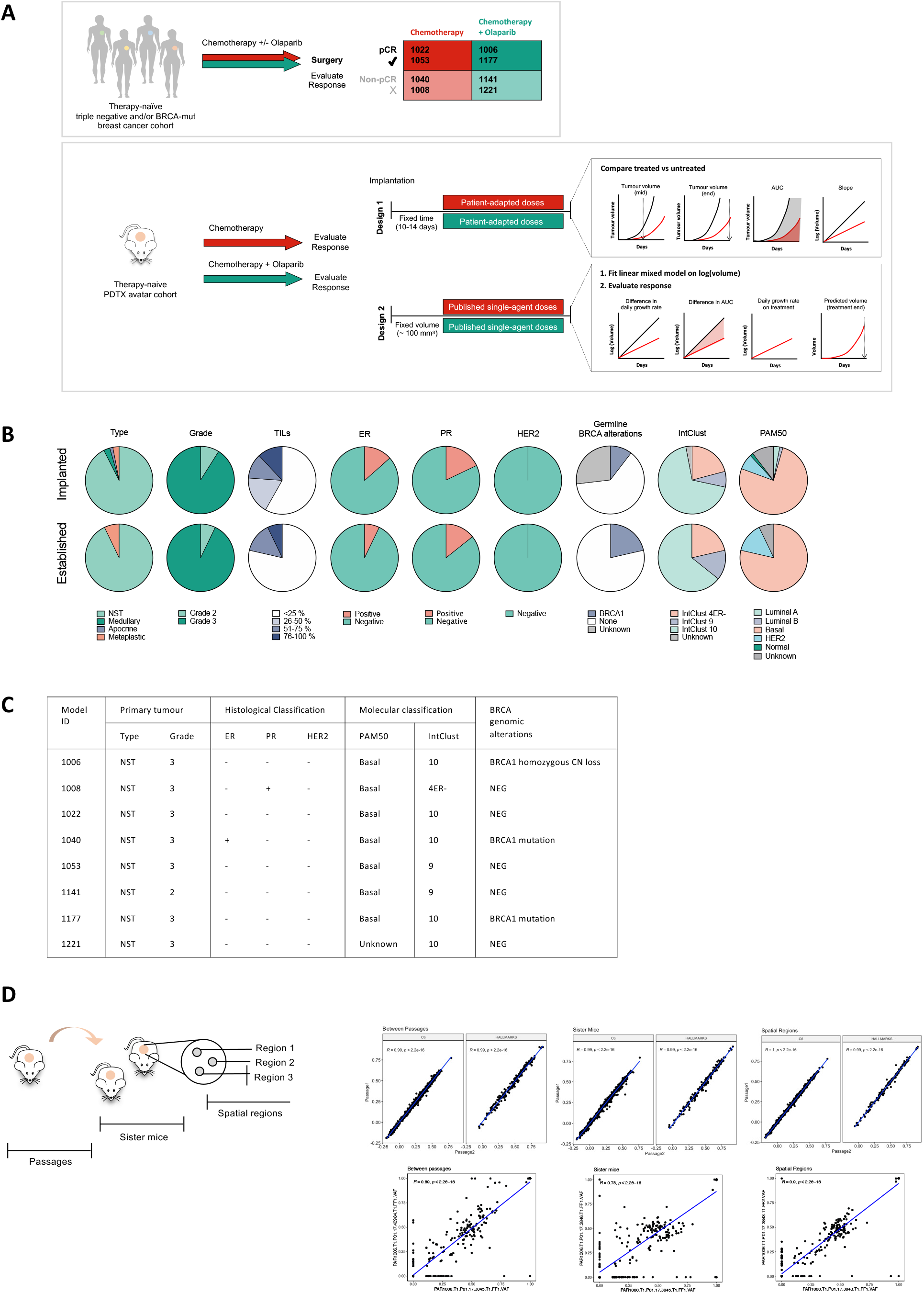
A cohort of triple-negative breast cancer (TNBC) patient-derived tumour xenografts (PDTXs) for co-clinical trials. A) Schematic of patient/PDTX experimental framework. B) Pie charts depicting the clinical features of the patient tumours from the PARTNER clinical trial, which were implanted and established as PDTX models. C) Clinical information of 8 PDTX models selected for the co-clinical trial. D) Correlation plots comparing enrichment scores from GSEA analysis using Hallmark and C6 gene sets for models 1006, 1040, 1022 and 1141 (top) and correlation plots comparing variant allele frequencies (VAF) mutations for model 1006 (bottom). Correlation was calculated between passages, sister mice and multiple regions of the same tumour. r value calculated using Spearman correlation.

PDTX models were subject to rigorous assessment by histology and immunohistochemistry and were shown to reflect the originating tumours (Supplementary Figure 1). Eight of the 14 established PDTX models were used for the study described here (unbiased selection based on availability of material). A summary of the clinical features is outlined in Figure 1C. The majority of tumours were TNBC, and two tumours weakly expressed either PR or ER. To minimise heterogeneity pre-randomisation, all PARTNER patients are assessed for basal phenotype using immunohistochemistry (ER; PR; HER2; CK5/6 and EGFR). All tumours had a basal phenotype and were classified into integrative clusters (IntClust) 4ER-, 9 or 10. Three tumours had genomic *BRCA1* alterations: homozygous copy number loss (model 1006), a pathogenic mutation (c.4327C>T, p.Arg1443Ter) with loss of heterozygosity (LOH) (model 1040), and a predicted benign mutation (c.2077G>A p.Asp693Asn) with LOH but associated with a BRCA-mutation-like mutational profile, indicating homologous recombination deficiency (model 1177).

RNA sequencing from four models (1006, 1040, 1022, 1141) was used to interrogate the impact of spatial heterogeneity on the expansion of PDTXs, by analysing distinct spatial regions of the same tumour, sister mice within a passage, and across multiple passages (Figure 1d, Supplementary Figure 2). Principal component analysis and hierarchical clustering of gene expression data revealed clear separations between models and samples from the same model clustered together. Gene set enrichment analysis was performed on each sample and we observed a high correlation between enrichment scores of both HALLMARK and oncogenic signature gene sets, C6, across spatial regions, sister mice and passages within each model. To further explore this at the mutational level, we performed whole exome sequencing on spatial regions, sister mice and across passages for one model (1006). This demonstrated a high correlation of the variant allele frequencies (VAF) of mutations between all levels. This supported our previous findings that the PDTX platform and experimental procedures were suitable for *in vivo* drug testing using sister mice as biological replicates and that the molecular features are stable across passages (19).

### A co-clinical trial reveals concordant PDTX-patient drug responses

To test the concordance of drug responses between matched patients and PDTXs, we developed a co-clinical trial platform. We tested two experimental trial designs and analytical frameworks to evaluate response (Figure 1A). For both, PDTX models were horizontally expanded into cohorts of daughter mice using n=5 per cohort following statistical power analysis. In design 1, to echo the PARTNER clinical trial regimen, treatment was administered 10-14 days post-engraftment. In design 2, treatment commenced at a fixed tumour volume (approximately 100 mm^3^) to account for model-specific growth dynamics; mice were assigned into cohorts using a stratified randomisation approach which aimed to evenly distribute initial tumour volumes. For both trial designs, mice were administered with the same treatment as the matched patient (named *avatar* mice) mirroring the patient treatment schedule and administration routes. In addition, the growth trajectory in the absence of treatment was monitored in 5 mice (named *untreated* mice). Patients and PDTXs were treated with chemotherapy (carboplatin and paclitaxel) +/- olaparib, herein referred to as CTO and CT trial arms respectively.

For design 1, avatar doses were adapted from the human doses in the PARTNER clinical trial relative to reference body mass ratios (human 60kg and mouse 20g). The human dose of carboplatin (AUC5) was converted to 0.16 mg fixed dose for mice. The human paclitaxel dose of 80 mg/m^2^ was converted to 0.04 mg but based on previous tolerability data on NSGs in our group, 0.07 mg was administered. The olaparib dose was converted from a 150 mg human dose to a 0.05 mg mouse dose. For design 2, drug doses were increased to 7mg/kg, 40mg/kg and 50mg/kg for paclitaxel, carboplatin and olaparib respectively after performing tolerability experiments in NSG mice. For both trial designs and to mirror the clinical treatment schedule, carboplatin was administered once every 3 weeks by intravenous injection, paclitaxel was administered intravenously once weekly and (when applicable) olaparib was administered by oral gavage on days 3-14 of each cycle. Four cycles of therapy were administered, with three weeks per cycle, to total 11 weeks (77 days) of treatment. While patients enrolled on the PARTNER clinical trial received three subsequent cycles of standard-of-care anthracycline chemotherapy, this was omitted for the co-clinical trial due to tolerability concerns in the mice.

To compare drug responses between trial designs, distinct analytical frameworks were adopted. For design 1, raw data was analysed and several parameters were used to compare the growth trajectory between the untreated and avatar mice: tumour volume after two treatment cycles (6 weeks, 42 days), tumour volume at the end of treatment (11 weeks, 77 days), the area under the curve (AUC) of the growth curve and the slope of growth curve following log2 transformation. For design 2, we adopted mathematical modelling to analyse the data. We used the assumption of exponential tumour growth (35) and fitted linear mixed models on the log-volume of the tumour on each day after the start of treatment for each PDTX model. We included an effect on treatment and a random slope for every mouse, allowing us to model the variability in growth rates observed between replicates. Four metrics were used to quantify the treatment effect: mean difference in daily growth rate and mean difference in AUC (both between treated and untreated groups), mean daily growth rate on treatment, and predicted tumour volume at treatment end. The mean difference in daily growth rate and AUC between the estimated marginal mean growth of the treated and the untreated groups were used to assess whether treatment caused any improvement in the mice over untreated controls, even in instances in which the tumour volume did not reduce. The mean daily growth rate for treated mice was used to identify whether tumour volume was changed from the start of treatment as an effect of the treatment. Predicted volume for each mouse at the end of treatment was considered to be the most similar metric to the pathological assessment performed in the patient, measuring the total tumour volume at the end of the treatment.

Both trial designs were tested in six PDTX models spanning both trial arms (CT and CTO), and patient responses (pathological complete response, pCR, and non-pCR). Trial design 2 was expanded to a further two PDTX models, and for those who displayed non-pCR, all possible residual cancer burden (RCB) scores are represented (I-III) (Figure 2).

**Figure 2:**
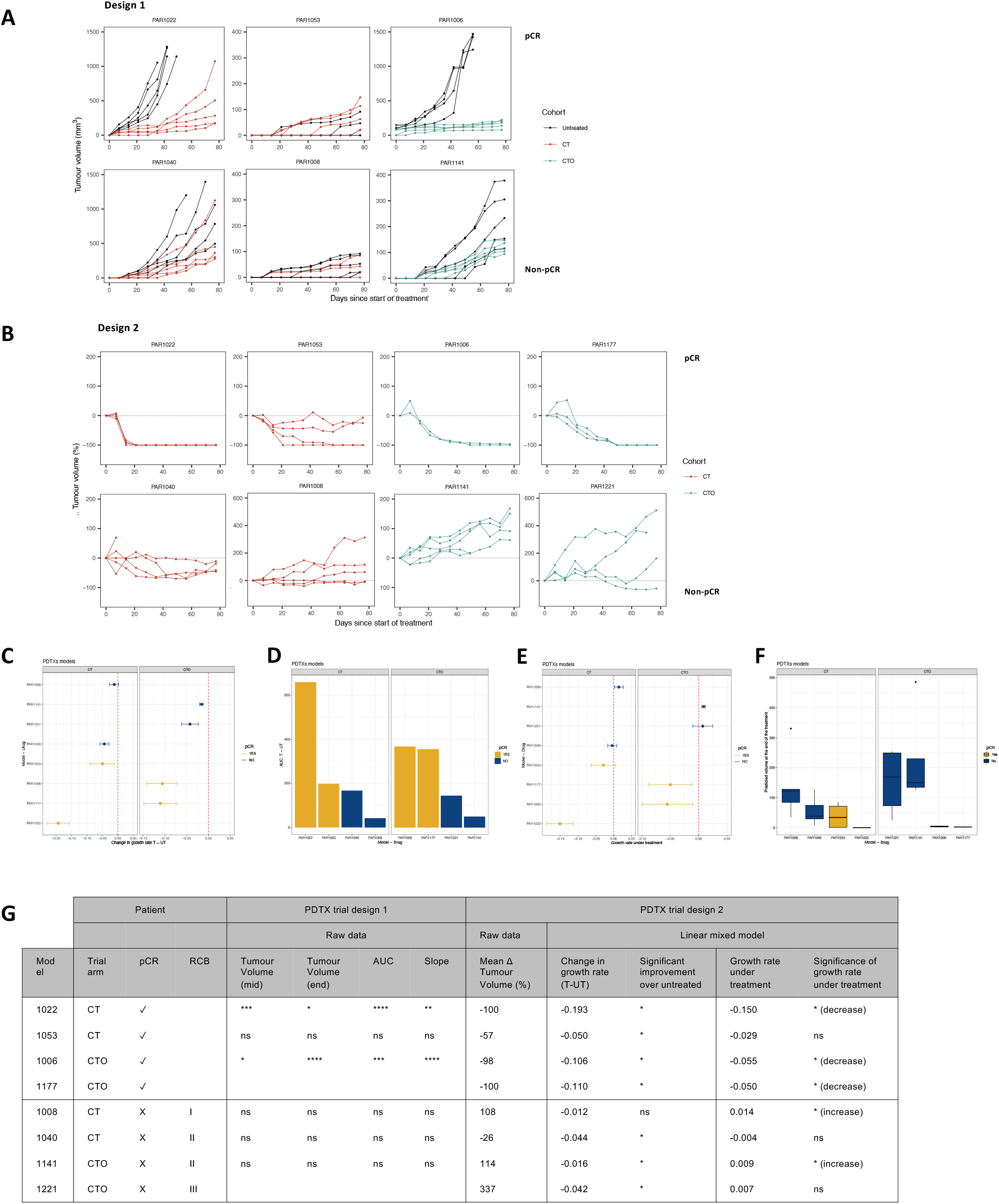
A co-clinical trial reveals concordant PDTX-patient drug responses *in vivo*. A) Growth curves of untreated and avatar cohorts treated using trial design 1 over 11 weeks of treatment. B) Percentage change (Δ) in tumour volume curves of avatar cohorts using trial design 2 over 11 weeks of treatment. C) Mean difference in daily growth rate between the treatment group and the untreated group, estimated with the interaction between growth rate and treatment group in the linear mixed model. D) Area under the curve between the estimated marginal mean growth of the treated and the untreated groups. E) Mean daily growth rates of mice in treated groups. F) Predicted volume for each mouse at the end of treatment. G) Results of the co-clinical trial (designs 1 and 2). For trial design 1, p values calculated using unpaired Welch’s t-test (unequal variance) comparing untreated to treated mice. * p<0.05, ** p<0.01, *** p<0.001, **** p<0.0001. ns: not significant. pCR: pathological complete response. AUC: area under the curve. CT: chemotherapy (paclitaxel and carboplatin). CTO: chemotherapy (paclitaxel and carboplatin) plus Olaparib.

The commencement of treatment at a fixed time point after implantation (trial design 1) was not suitable for models with slower proliferation rates, e.g. 1008 and 1053. For these models, several untreated mice did not display measurable tumours after the trial ended, which posed challenges when determining effects of treatment on tumour growth. In the case of model 1053, this donor patient achieved a clinical pCR, but no significant differences were observed between the untreated and avatar mice by all parameters analysed.

We also asked whether the low doses used in design 1 could be confounding the results. PDTX tumour growth was detected even in cases in which the corresponding patient achieved pCR, e.g. models 1006 and 1022. While this could be attributed to the additional cycles of anthracycline chemotherapy the patients received, the drug doses used in design 2 reduced PDTX tumours to unmeasurable levels in these models, representative of clinical pCR. In addition, commencing treatment at a fixed tumour volume accounted for model-specific growth dynamics, which along with higher administered doses, resulted in tumour regression.

Data from trial design 2 revealed high concordance between patient and PDTX drug responses. While these data revealed significant improvements in treated mice compared to untreated controls in 7/8 models tested, significant tumour regression (indicated by negative growth rate under treatment) was observed only in models generated from a matched patient that achieved pCR. Likewise, only models from patients which achieved a clinical pCR had a predicted tumour volume of 0 at treatment end. We observed high variability between biological replicates for model 1053 but, across the cohort of mice, change in growth rate (T-UT), daily growth rate under treatment and tumour volume at treatment end were all lower than all PDTX models from which the matched patient had non-pCR.

These data comprise the first systematic assessment of concordant drug responses in a neoadjuvant treatment naive breast cancer setting using two distinct trial designs. We demonstrated an accurate recapitulation of clinical drug responses between matched patient-PDTXs in eight models tested using several analytical metrics. These data elegantly exemplify the power of a co-clinical trial platform in predicting patient’s drug responses setting strong grounds for exploiting the use of PDTXs to anticipate the most efficacious treatment strategies in patients that are not responding to standard-of-care or that later relapse.

### Using PDTXs to identify alternative efficacious treatment strategies and to interrogate patient- and treatment-specific tumour evolution

PDTX models have potential utility as tools in anticipatory cancer medicine to identify efficacious treatment strategies for patients and to model patient- and drug-specific tumour evolution (Figure 3A).

**Figure 3:**
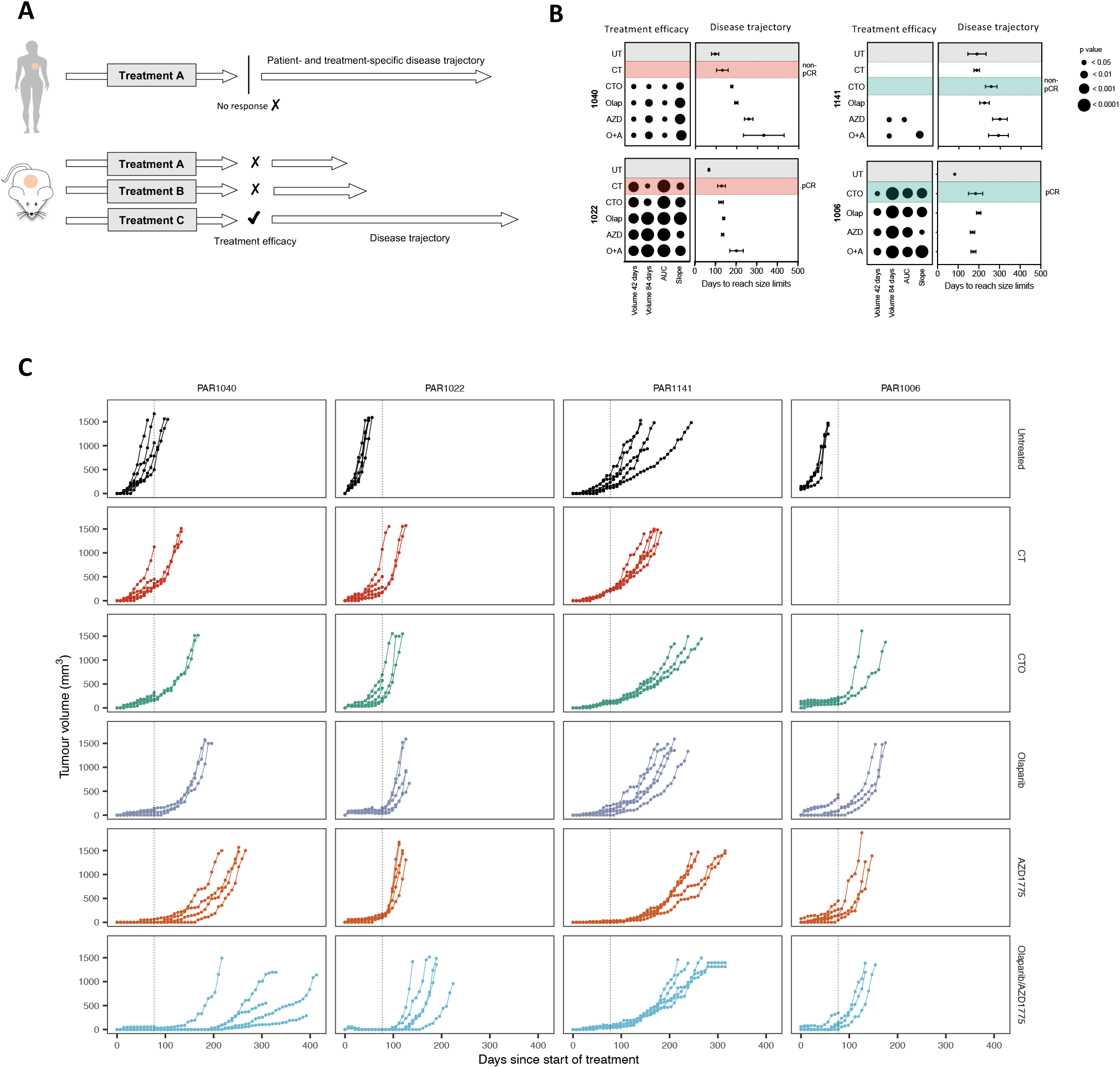
Using PDTXs to identify alternative efficacious treatment strategies and to interrogate patient- and treatment-specific tumour evolution. A) Schematic of experimental design. B) Treatment efficacy and disease trajectory of four PDTX models treated with avatar (highlighted red/green) and alternative treatment strategies for 11 weeks. Disease trajectory is defined as days to reach size limits of 1500mm^3^ from start of treatment. UT: untreated. CT: chemotherapy. CTO: chemotherapy (paclitaxel and carboplatin) plus Olaparib. Olap: Olaparib monotherapy. AZD: AZD1775. O+A: Olaparib and AZD1775 combination. C) Tumour growth curves of four models: untreated or treated with avatar or alternative strategies. Dotted line indicates end of treatment.

Four distinct TNBC PDTX models were treated *in vivo* with the same (avatar) or the alternative clinical trial arm to what the donor patient received, and also with other clinically-relevant therapies for 11 weeks (Figure 3B-C). After treatment withdrawal, mice were left untreated to identify tumour relapse. This enabled us to explore both drug efficacy on treatment and the long-term evolutionary dynamics for every patient-treatment pair. We observed different dynamics of relapse in a patient- and treatment-specific manner. These results were particularly pronounced and of potential clinical relevance for models 1040 and 1141. For 1040 neither the patient nor the PDTX avatar responded to CT, while all alternative treatment strategies were successful to a similar degree during the treatment course. However, once treatment ceased the growth trajectories displayed marked differences: following CTO treatment, tumours reached 1500mm^3^ in a mean time of 176 days from implantation and following combination treatment with olaparib and AZD1775 tumours reach 1500mm^3^ in a mean time of 331 days. Model 1141 was largely resistant to treatment *in vivo* but responded to AZD1775 both as a monotherapy and in combination with olaparib. In stark contrast, model 1022 was highly sensitive to all compounds tested. These data support the potential use of a PDTX drug screening platform as a tool for precision cancer medicine to identify patient’s drug responses and evolutionary trajectories.

### Using a PDTX drug screening platform to interrogate mechanisms of drug response to olaparib

The *in vivo* drug screening platform revealed that models 1006 and 1040 (both with genomic BRCA1 alterations) were highly responsive to olaparib (both monotherapy and in combination with chemotherapy or AZD1775). Model 1022 was also highly responsive to olaparib, and upon further investigation it was revealed to be BRCA1-null at the transcriptomic and protein level (Supplementary figure 3B-C). As expected, model 1141 did not respond to olaparib (homologous recombination proficient). However, despite this high responsiveness in models 1006, 1022 and 1040, rapid tumour regrowth was observed upon drug withdrawal. This indicated that a population of viable cells persisted and/or continued to proliferate on treatment, subsequently leading to relapsed disease.

While numerous PARPi resistance mechanisms have been described (33), preclinical resistance studies typically have limited relevance to the clinical setting. Here we leveraged our unique platform using therapy-naïve PDTXs and clinically-relevant treatment schedules to identify mechanisms which may be more closely applicable to clinical tumour evolution. Although we did not observe reversion of the BRCA1 mutation in model 1040 (Supplementary figure 3D), we selected model 1006 to perform a deep-dive analysis so as to avoid the chance of PARPi resistance being caused by secondary mutations and enable the identification of other clinically-relevant new mechanisms directing cancer evolution in TNBCs. Model 1006 has both a germline and somatic BRCA1 copy number loss and does not express BRCA1 at the RNA or protein level (Supplementary Figure 3A-C). Although in the PARTNER clinical trial, olaparib is used in combination with chemotherapy, for our purpose we analysed PDTX samples treated with olaparib monotherapy to identify PARPi-specific mechanisms. Tumours from biological replicates were harvested at two time points to temporally resolve the mechanisms of drug response dynamics, including relapse: immediately following treatment (denoted as *treated)* and at size limits after drug withdrawal *(post-treated)*, modelling residual and relapsed disease respectively (Figure 4A).

**Figure 4:**
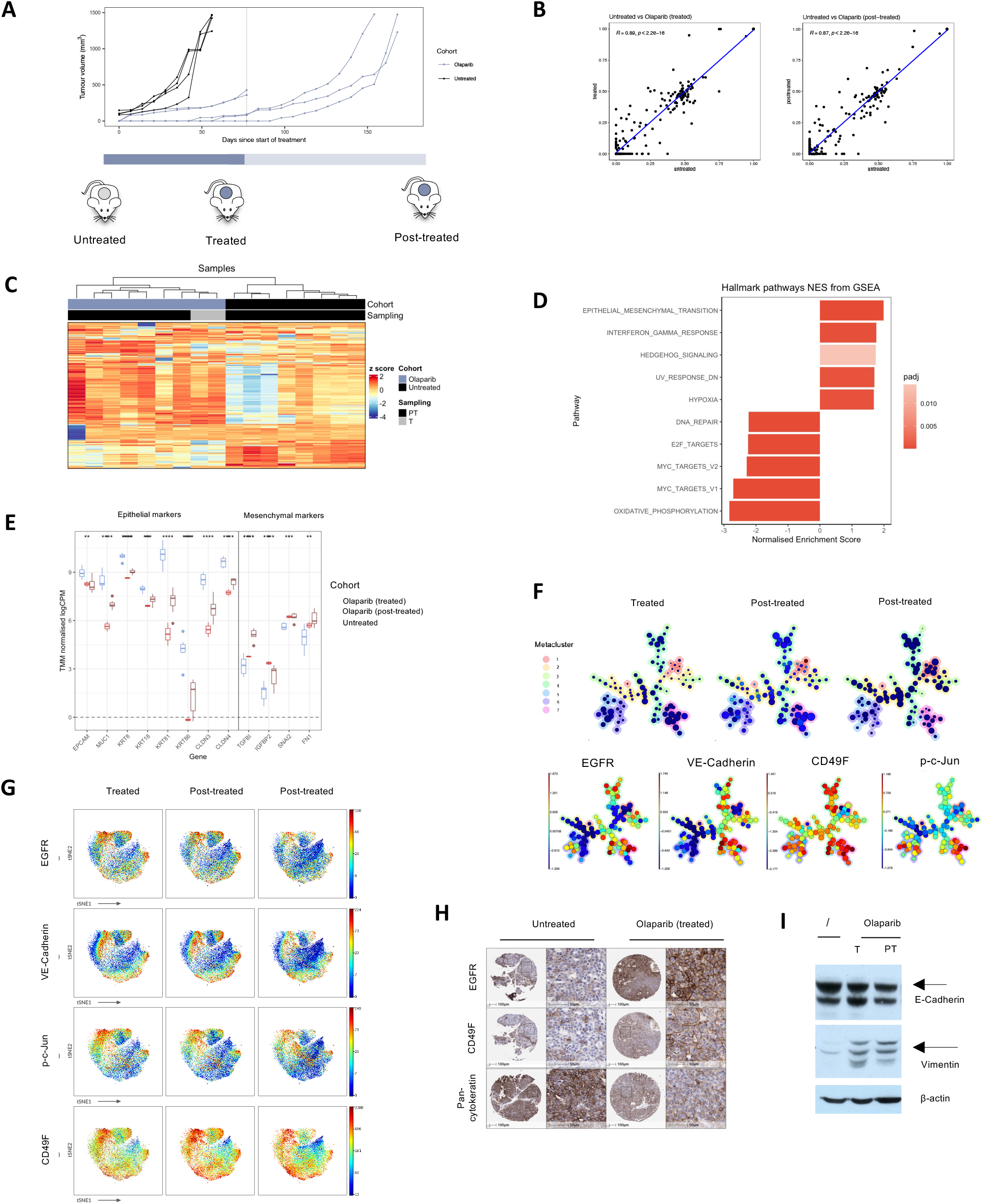
Olaparib induces permanent non-genetic phenotypic changes in TNBC. A) *In vivo* growth curves of model 1006 treated with olaparib (purple) or untreated controls (black). Dotted line indicates end of treatment. B) Correlation plots comparing mean variant allele frequencies (VAF) of mutations between untreated and treated (left) or untreated and post-treated (right) samples. r value calculated using Spearman correlation. C) Heatmap showing z score (scaled by row) of the top 250 strong and variable genes in PDTX samples. Clustering analysis performed using Euclidean distances. Columns indicate samples. T: Treated. PT: Post-treated. D) Top 10 significant gene sets by normalised enrichment score, identified by gene set enrichment analysis using the HALLMARK gene sets between untreated and post-treated samples. E) Normalised gene expression of epithelial and mesenchymal markers. All genes were found to be differentially expressed between untreated and post-treated samples. F) viSNE plots treated and post-treated samples (columns) displaying the expression of cell type markers (rows) measured by CyTOF mass cytometry. G) Minimum spanning tree of representative samples (top). Inner circle density represents event counts per cluster and outer circle colour indicates metacluster. Expression of markers in each cluster is displayed in the bottom panel. H) Immunohistochemistry for key markers in untreated and treated samples. I) Western blot for E-cadherin and vimentin in untreated, treated (T) and post-treated (PT) samples.

To interrogate the mechanisms by which tumours survived upon treatment, we performed a deep molecular characterisation of treated and post-treated tumours using several analytical modalities. Genomic analysis using multi-region whole exome sequencing (Supplementary Figure 4) failed to identify any mutations which could fully explain the survival of cells on treatment. Filtering non-synonymous SNVs, frameshift indels and stop-gain mutations, we observed a high correlation between the VAFs detected in untreated, treated and post-treated tumours (Figure 4B). Hierarchical clustering of SNV VAFs revealed no clear separation between untreated, treated and post-treated tumours (Supplementary Figure 4C). We then sought to identify ‘emergent’ and ‘depleted’ mutations. Emergent mutations were defined as those present in no untreated samples, but in at least one treated or post-treated sample; conversely depleted mutations were those present in two or more independent untreated mice, but no treated or post-treated samples. 13 depleted mutations and 28 emergent mutations were identified (Supplementary Figure 4D). While most were classified as predicted passenger mutations, we identified a depleted mutation in GATA2 and an emergent mutation in TRRAP, both predicted driver mutations (36, 37). However, since these drivers were detected in just two untreated mice and one post-treated mouse respectively, it was deemed insufficient to fully explain natural selection of a genetic clone through Darwinian evolution under a treatment regime during this time-frame *in vivo*.

We next analysed gene expression by bulk RNA-sequencing. We observed minor yet significant changes in gene expression of known PARPi resistance markers related to DDR rewiring, including BRCA2, RAD51, PALB2 (Supplementary Figure 5A). Albeit potentially involved in driving resistance, here we opted to instead adopt an unbiased and holistic data-driven approach. Using the top 250 strong and variable genes identified in bulk RNA-sequencing, samples were clustered by Euclidean distance (Figure 4C). Treated and post-treated tumours clustered together, and separately from untreated tumours. To our surprise, this demonstrated a global transcriptomic shift upon treatment with olaparib which persisted (irreversible) when treatment was removed, even when left 3 months off treatment. Gene set enrichment analysis was performed between untreated and post-treated tumours (Figure 4D, Supplementary Figure 5B) which demonstrated a significant enrichment of the HALLMARK ‘Epithelial to Mesenchymal Transition’ gene set. Differentially expressed genes were then identified between untreated and post-treated tumours; post-treated tumours exhibited up- and down-regulation of several mesenchymal and epithelial genes respectively (Figure 5E).

**Figure 5:**
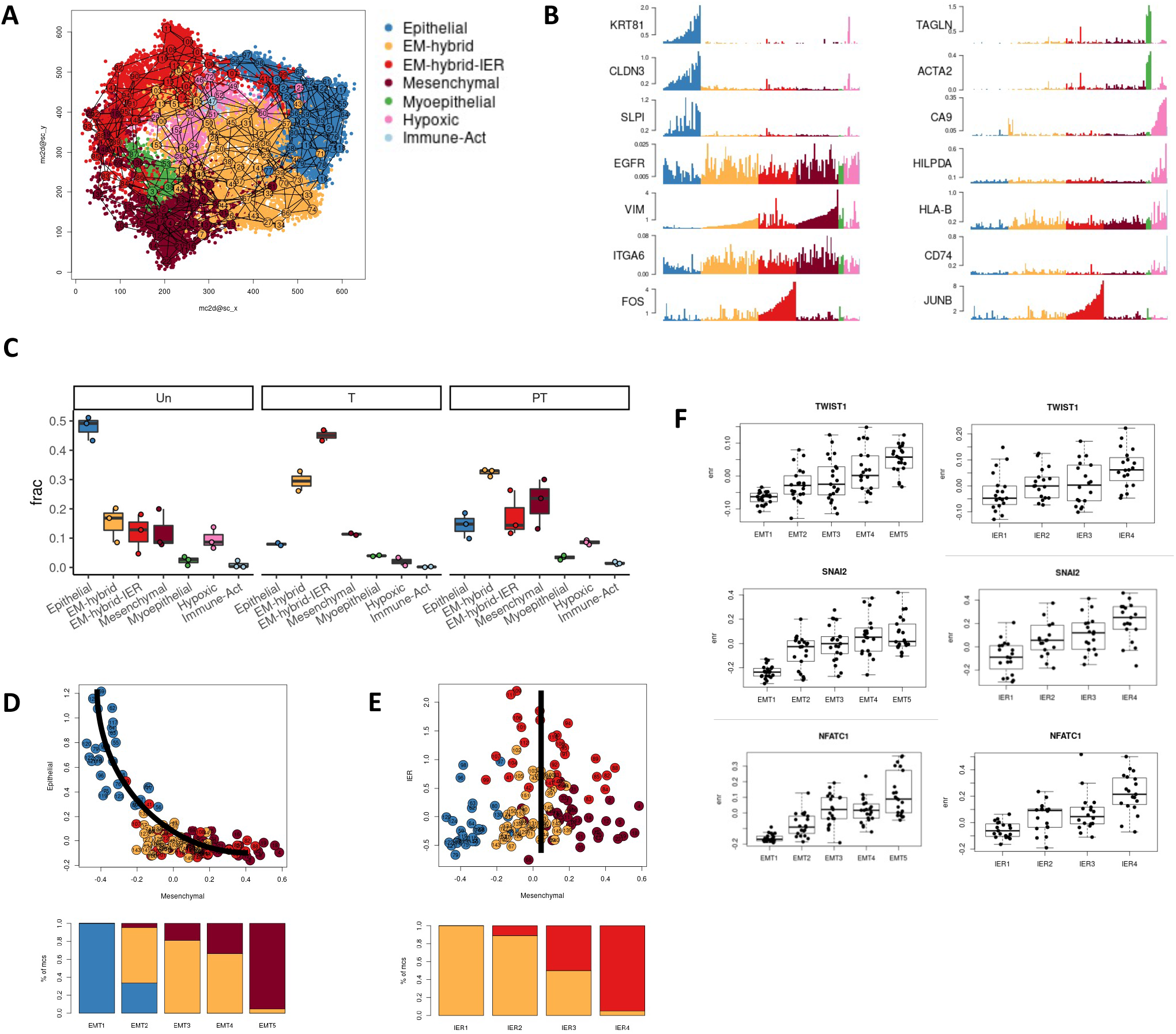
Single-cell RNA-sequencing reveals transient and permanent changes in phenotypic populations upon treatment driven by parallel transcriptional programmes, towards a mesenchymal, stem-like state. A) Single-cell RNA-sequencing data of all cells analysed post-QC from model 1006. Colour indicates cells states (groups of metacells). B) Average gene expression of key markers across metacells. C) Fraction of cells from untreated, treated (T) and post-treated (PT) samples to reside cell states. D) Epithelial and Mesenchymal scores for individual metacells (top). Line indicates the gradient captured using stratification. Metacell composition of each strata (bottom) using mesenchymal minus epithelial scores to stratify metacells into 5 groups (EMT1-5). E) Mesenchymal and IER scores for individual metacells (top). Line indicates the gradient captured using stratification. Metacell composition of each strata (bottom) using IER score to stratify metacells into 4 groups (IER1-4). F) Mean enrichment across strata (EMT1-5 left, IER1-4 right) for genes of interest.

Model 1022 (transcriptomic BRCA1-null) demonstrated a significant and dramatic increase in BRCA1 expression in tumours treated with olaparib using the same experimental design (Supplementary Figure 5C-D). After treatment with olaparib, 5/5 mice demonstrated significantly higher gene expression and indeed the most highly significant differentially expressed gene between untreated and post-treated tumours was BRCA1 (logFC 9.06, p-value 4.99 e-06), indicating an epigenetic reversion of gene suppression. These data suggest that our platform enables the identification of known single-biomarkers of resistance to PARPi. Crucially, these data also identify new molecular dynamics involved in olaparib-dependent drug responses, and suggests this is driven by a global non-reversible transcriptomic change.

To validate these findings at the phenotypic level, we adopted our previously described time-of-flight mass cytometry (CyTOF) approach (20) in dissociated cancer cells (Figure 4F-G, Supplementary Figure 6). Data were analysed using Cytobank (38). Treated and post-treated tumours demonstrated similar phenotypic profiles by viSNE and FlowSOM clustering, supporting the concept that the phenotype observed immediately after treatment was largely stable following drug withdrawal. We did observe minor changes, such as increased metacluster 4 and decreased metacluster 7 in post-treated samples compared to treated; both metaclusters highly expressed CD49F, EGFR and VE-Cadherin, but metacluster 7 exhibited high expression of p-c-Jun and other signalling proteins. In addition, we observed a decrease in metacluster 5 (epithelial phenotype), suggesting a shift towards a more mesenchymal phenotype in post-treated compared to treated samples. Mass cytomery was then performed on untreated PDTX tumours from model 1006 independently propagated, which revealed a homogenous low expression of EGFR and VE-Cadherin. These data suggest that treatment with olaparib induces a progressive switch towards mesenchymal-like cell states, by increasing populations enriched in these markers, which continue to increase upon drug withdrawal. These data were further validated at the tissue level using immunohistochemistry (Figure 4H), which demonstrated an increase in expression of EGFR and CD49F and a decrease in expression of cytokeratins in treated compared to untreated trial samples. Likewise, a western blot of homogenised tissue revealed a stepwise decrease in E-Cadherin and increase in Vimentin, epithelial and mesenchymal markers respectively (Figure 4I). Taken together, these data demonstrate at both the expression and protein level that non-genetic phenotypic changes occur upon treatment with olaparib; persistent and emergent populations are enriched for mesenchymal markers and phenotypic changes remain irreversible after drug withdrawal, with transient high expression of signalling markers.

To further interrogate the molecular mechanisms underlying these changes, single cell RNA-sequencing was employed. 21,894 cells from eight samples (at least two biological replicates per condition) were available for analysis after applying quality control filters. *Metacell* analysis (39) revealed 7 groups of metacells, or cell states (Figure 5A-B, Supplementary Figure 7A). Minor cell states expressed high levels of myoepithelial (e.g. TAGLN, ACTA2), hypoxic (CA9, HILPDA) and Immune Activation (HLA-B, CD74) markers. The remaining metacells were classified into cell states based on expression of epithelial (e.g. KRT81, CLDN3, SLPI) and mesenchymal markers (e.g. VIM, ITGA6, EGFR). Metacells were divided into Epithelial, Mesenchymal and EM-hybrid states, and a subset of EM-hybrid metacells expressed high levels of immediate early response (IER) genes (e.g. FOS, JUNB), which we denoted as EM-hybrid-IER.

The majority of cells in untreated samples exhibited an epithelial phenotype, consistent with bulk RNA-sequencing and protein-level data (Figure 5C). Immediately after treatment, there was a dramatic decrease in epithelial and increases in EM-hybrid and EM-hybrid-IER cells. After treatment withdrawal, there remained high levels of EM-hybrid cells (but fewer with IER expression) and an increase in mesenchymal cells. We also observed a small increase in epithelial cells, but this remained dramatically lower than in untreated controls.

These data revealed two key phenotypic changes induced by olaparib treatment, with both transient and permanent dynamics. In line with our previous data, olaparib caused phenotypic changes involving sequential transitions from an epithelial to an EM-hybrid cell state on treatment, and to a fixed mesenchymal cell state in relapse samples. While we did observe a small degree of reversion to an epithelial phenotype, the overwhelming majority of post-treated cells remained EM-hybrid or mesenchymal. In addition, this single cell analysis revealed that olaparib induced transient high expression of IER genes in cells residing in a stressed cell state, supporting the mass cytometry data. Crucially, these changes were highly consistent between both biological and technical replicates (Supplementary Figure 7C).

To further elucidate the drivers of this, we calculated three gene expression scores (Epithelial, Mesenchymal, IER) by averaging gene enrichment of the top 50 genes correlated with KRT81, VIM and JUNB respectively (Supplementary Figure 8). Using expression data from Epithelial, EM-hybrid and mesenchymal metacells, each score was correlated with the expression of known transcription factors (TFs) (40). As expected, TFs which correlated with the epithelial score were negatively correlated with the mesenchymal score, and vice versa. Several TFs known to be associated with epithelial and mesenchymal phenotypes were indeed correlated with the respective scores, such as ELF5 (epithelial) (41) and TWIST2 (mesenchymal) (42). A subset of TFs were found to be correlated with both the mesenchymal and IER expression scores, including EMT master regulator SNAI2 (43). This led us to hypothesise that the two key phenotypic changes observed in response to olaparib may be interrelated and that the transient overexpression of IER genes may drive, or be associated with, the permanent gene expression changes upon treatment.

Epithelial and mesenchymal scores were plotted for individual metacells which (in line with several recent reports) demonstrated that EMT is a continuum, rather than discrete cell states (Figure 5D) (44–47). Epithelial, mesenchymal and EM-hybrid metacells were stratified into five groups (EMT1-5). Mean gene enrichment of TFs across the EMT1-5 strata was used to infer TFs which may be driving different stages of the EMT trajectory (Supplementary Figure 9). For some TFs, gene enrichment increased or decreased step-wise along the continuum, for example TWIST1 and ELF5 respectively. Others appeared to be early-acting, establishing a hybrid EM cell state e.g. SNAI2, ATF5. Conversely, other TFs were late-acting, transitioning cells from EM-hybrid to a fully mesenchymal phenotype, such as TWIST2, IRX3. Adopting a similar approach for the IER trajectory (Figure 5E), EM-hybrid and EM-hybrid-IER metacells were stratified into four groups (IER1-4). Again, mean gene enrichment of TFs was plotted across the IER1-4 strata (Supplementary figure 9), which identified several TFs which correlated with acquisition of an IER phenotype, including TWIST1, SNAI2 and NFATC1.

The observation that several TFs overlapped between the EMT and IER trajectories (Figure 5F) supported our hypothesis that the transient and permanent phenotypic changes were interrelated. These data suggest that transcriptional programmes associated with an IER trigger a cascade of phenotypic state dynamics which occur independently or in parallel/interplaying with clonal evolution.

These data demonstrate the value of leveraging the PDTX platform to reveal patient-specific evolutionary behaviours and to uncover unidentified molecular mechanisms and phenotypic dynamics driving relapse.

### Identifying new therapeutic vulnerabilities using patient-specific evolutionary insights

We next tested the functional implications of *in vivo* drug screening in four PDTX models (Figure 6A). We performed high-throughput drug screens on dissociated PDTX cells which had previously been pre-treated and subject to drug withdrawal (as in Figure 3). This revealed both patient- and drug-specific functional changes, as demonstrated by area under the curve (AUC) values.

**Figure 6:**
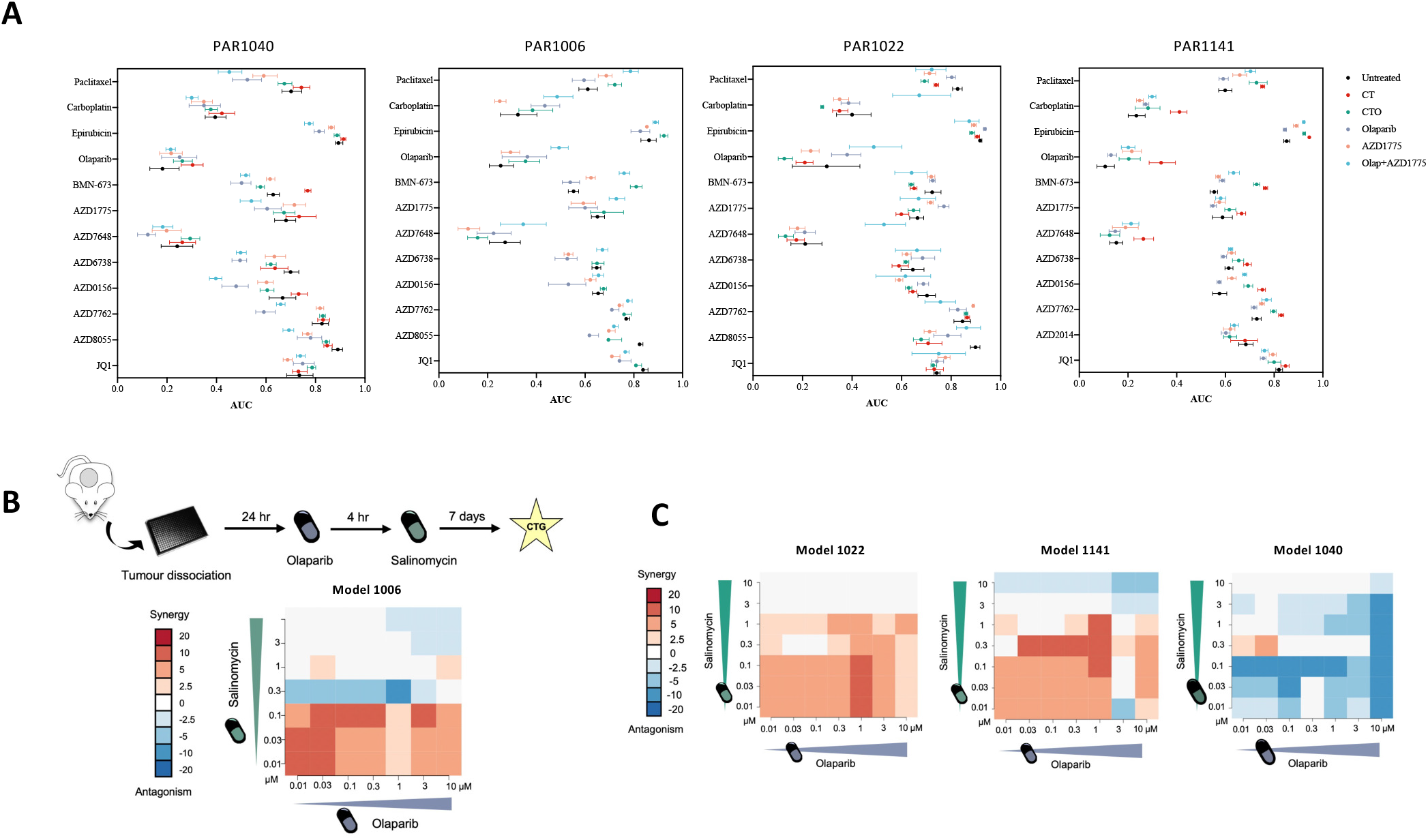
Identifying new therapeutic vulnerabilities using patient-specific evolutionary insights. A) *Ex vivo* high-throughput drug screening of PDTX tumours from four models, following *in vivo* treatment with avatar and alternative treatment strategies. Plots display area under the curve (AUC) values. B) Experimental design and heatmap of synergy between olaparib and salinomycin when treated in combination in dissociated cells from model 1006. Synergy calculated using Bliss model. C) Heatmap of synergy score between olaparib and salinomycin for models 1022, 1141 and 1040.

Using the evolutionary insight gained from the deep-dive analysis, we set out to identify efficacious patient-specific therapeutic strategies which could involve sequential or upfront combination therapies for model 1006. A potassium ionophore, salinomycin, has been identified to selectively target cells which have been transitioned through an EMT (48). We postulated that, given the increase of a mesenchymal phenotype after *in vivo* olaparib treatment, olaparib followed by salinomycin could be an efficacious sequential therapy. An *ex vivo* combination drug screen was performed on dissociated cells from model 1006, first dosing with olaparib and then salinomycin 4 hours later (Figure 6B). Using the Bliss model (49), significant synergy was identified between the two compounds, supporting the high efficacy of the use of these drugs in sequence. Furthermore, this experiment supports the concept that olaparib induces transitions to other cell states (phenotypic switching) as opposed to an enrichment of a pre-existing population. A selective killing of different populations by the two compounds would lead to an additive, rather than the observed synergistic effect. From this, we propose that olaparib induces cell plasticity to a phenotypic state (mesenchymal) that is selectively killed by salinomycin.

To test whether this phenomenon was specific to olaparib or independent of the therapeutic pressure, we tested the combination of carboplatin and salinomycin. No synergy was observed (Supplementary Figure 10), indicating that the mechanism is drug-specific. The experiment was then extended to other PDTX models and remarkably 3/4 models demonstrated synergy between olaparib and salinomycin (Figure 6C), suggesting this phenotypic behaviour is a common phenomenon. Strikingly, when olaparib and salinomycin were used in upfront combination (rather than sequential) the degree of synergy was lower, further supporting the plasticity hypothesis that olaparib primes cells for effective salinomycin killing.

Taken together, these data demonstrate the value of translating evolutionary and phenotypic insights from the PDTX platform to identify personalised anti-cancer therapeutic strategies based on patient-specific behaviours, highly predictive of patient drug responses. It also suggests that salinomycin could be a potential avenue for therapeutic intervention for relapsed TNBC following treatment with olaparib.

## DISCUSSION

Breast cancers are highly complex cellular entities, exhibiting both inter- and intra-tumour heterogeneity. TNBCs remain the subgroup with the worst prognosis and, while DDR agents offer new avenues for therapeutic intervention, drug resistance and disease relapse remain obstacles in the treatment of the disease. To fully understand the mechanisms by which tumours evade therapy and evolutionary principles that drive its progression, it is essential to leverage preclinical models which accurately recapitulate the heterogeneity of human breast cancer and adopt holistic tools which resolve such complexity.

We developed and optimised a co-clinical trial platform alongside an ongoing clinical trial using treatment-naïve TNBC PDTXs, and demonstrated concordance between drug responses of patients and their corresponding PDTX models using clinical endpoints. Treatment-naïve PDTXs mirror matched patient’s drug responses, establishing very strong foundations for the use of drug response information from co-clinical trial platforms to be adopted in precision medicine approaches. Using PDTX *in vivo* preclinical trials, we expanded the number of compounds a given patient’s tumour received and modelled patient- and drug-specific tumour evolution. This shows that the platform can be coupled and expanded to interrogate additional clinically-relevant therapeutic approaches in parallel, and be used as a predictive or anticipatory tool in patients who are not responding to treatment or that relapse. We then exploited this platform to interrogate mechanisms of drug response to olaparib. Using two analytical modalities at a single cell resolution (mass cytometry and RNA-sequencing) we identified a novel non-genetic mechanism of drug response to olaparib. This is driven by phenotypic cell dynamics (or cell plasticity), directed by transcription factor-mediated gene expression transitions that move cells from an epithelial towards stable mesenchymal states and that can be selectively targeted with salinomycin.

While we are not the first to report concordant drug responses in breast cancer (11, 12, 50), we present a robust approach to assess the extent to which PDTXs recapitulate clinical drug responses in a controlled clinical trial setting. The co-clinical trial and PDTX drug screening platform have wide-ranging implications for the future use of PDTXs in both anticipatory cancer medicine and the study of cancer evolution. It offers the opportunity to prospectively anticipate (i.e., predict) alternative efficacious treatment strategies for patients, particularly in the relapsed or recurrent setting. The timeframes reported here would be suitable to evaluate treatment options for relapsed tumours if PDTXs were generated from a patient’s sample upon initial disease presentation (including tumour biopsies). It also allows one to study the impact of that treatment on the tumour’s ability to evolve and interrogate parallel evolutionary trajectories upon different therapeutic pressure. While expected, an interesting observation was the dramatically different dynamics of relapse in a patient- and treatment-specific manner. This is critical data when designing treatment schedules as, even when treatments share good responses, each could follow distinct, synergistic or synthetically lethal evolutionary paths, highlighting the potential to leverage the concept of therapeutic steering towards chronicity based on evolutionary knowledge (51). The PDTC drug screening platform previously pioneered by our group (19) was further integrated into this work and offers the potential to screen large numbers of novel and clinical compounds in a high-throughput manner. It also confirmed that non-genetic cell plasticity-driven changes upon olaparib treatment had functional (treatment response) implications.

Using clinically relevant time frames and parallel treatment schedules to the patient, this study had major benefits over traditional drug resistance studies, more closely mimicking the clinical scenario. The mechanism of response to olaparib described here is akin to previous reports of phenotypic switching in response to treatment (52–55). A striking observation was the permanent nature of the phenotypic change, contrasting with mechanisms such as drug-tolerant persister cells (56), but more in line with stable non-genetic drug-resistance as described in (54, 55). Our single-cell RNA-sequencing data suggests that upon treatment cells adopt an EM-hybrid cell state enriched for IER genes, but upon drug withdrawal cells continue to progress towards a fully transitioned and stable mesenchymal phenotype. Previous reports have demonstrated that EM-hybrid cells exhibit higher metastatic potential (45) and tumorigenicity (57) than their epithelial or mesenchymal counterparts. Our data leads to the compelling hypothesis that drug withdrawal may drive cells further towards a mesenchymal cell state which, although traditionally associated with a more aggressive phenotype than epithelial cells, has less metastatic potential and tumorigenicity than hybrid cells that emerge on treatment. Using the single cell data to derive EMT and IER trajectories, we identified that overlapping TFs (including EMT master regulator SNAI2) may be implicated in directing cell state transitions, highlighting the potential of identifying plasticity-driver vulnerabilities. The timeframe of this approach may not have been long enough to identify natural selection of advantageous mutations. However plausible, the data presented here fits with the new compelling evidence supporting non-genetic evolutionary cancer drivers in the early stages of adaptation to treatment. As such, we hypothesise that targeting the molecular drivers that enable cancer evolution instead of/in addition to targeting the evolutionary endpoint, especially in the curative/early setting, will provide the recipe for higher clinical benefit. Outstanding questions relate to the dependency and sequence of this TF activation. One explanation could be that IER gene expression on treatment activates transcriptional programmes associated with EMT, priming cells to become mesenchymal even once IER gene expression reduces upon drug withdrawal. Alternatively, both IER and EMT processes may be driven by the same transcriptional master regulators induced by olaparib treatment; IER genes may be able to reduce back to baseline upon drug withdrawal but EMT, once established, continues along its trajectory towards a stable mesenchymal phenotype.

Our observation of synergy between salinomycin and olaparib further supports targeting phenotypic changes as a powerful therapeutic strategy. Indeed, salinomycin is gaining traction for treatment of several cancer types owing to its high selectivity towards cancer stems cells (48).

While mimicking the treatment schedules of the clinical trial compounds (carboplatin, paclitaxel and olaparib) in PDTXs in the co-clinical trials, patients received three subsequent cycles of standard-of-care anthracycline chemotherapy which was omitted. However, we argue that this work provides a strong foundation and a preliminary experimental framework to further expand into larger patient cohorts and a wider range of clinical compounds and clinical scenarios. Ultimately, a clinical trial with an avatar arm is needed to consolidate the use of this data to benefit cancer patients.

The mechanism of drug response described here supports mechanisms described previously and provides an interesting avenue for further exploration. A major limitation with our model system is that longitudinal sample collection provides only a snapshot in time, making it challenging to delineate between phenotypic switching and enrichment of pre-existing populations. The synergy observed between salinomycin and olaparib supports the hypothesis of phenotypic switching, but a barcoding approach for lineage tracing coupled with single-cell RNA-sequencing technologies would provide more granular information. Furthermore, we propose that salinomycin could be an interesting drug candidate for TNBC (either relapsed or in combination as a first-line therapy) and this warrants further exploration.

We describe a unique and robust platform to study drug responses and cancer evolution in a clinically relevant manner. This study has profound implications for the future use of PDTXs as both anticipatory tools for subsequent treatments in clinical medicine and a preclinical approach to high-resolution mapping of drug response dynamics.

## METHODS

### Generation and maintenance of PDTXs

Breast cancer patient-derived tumour xenografts (PDTX) were established and passaged as described in (19). Patient primary breast cancer biopsy samples were collected at diagnosis and immediately transported to the animal facility. Patient tissue samples were embedded in Matrigel and implanted subcutaneously into ~2 female severe immunocompromised NOD.Cg-Prkdcscid Il2rgtm1Wjl/SzJ (NSG) mice. PDTXs were serially implanted into multiple hosts to allow for *in vivo* expansion of each model. Xenograft samples were flash frozen in liquid nitrogen, cryopreserved in freezing media (FBS/10% DMSO) and fixed in 10% neutral buffered formalin at each passage from each mouse. Regular genotyping and immunohistochemistry quality control was performed routinely to confirm the propagation of human breast cancer tissue matching the originating patient-derived sample. All use of human samples and xenograft generation is covered by the appropriate human ethics framework in the UK, and all animal work is performed under the Home Office regulatory framework (project licence number: P1266F82E). The research was done with the appropriate ethical approval and informed consent was obtained from all patients. PARTNER samples were collected and held under the PARTNER ethics: IRAS project ID: 178681, Haydock Research Ethics Committee. REC reference number for PARTNER: 15/NW/0926. REC reference number for PBCP: 18/EE/0251.

### *In vivo* preclinical drug studies

Fresh or cryopreserved tumour fragments from a single mouse were passaged into a cohort of NSG mice. At the appropriate point (either a defined time after implantation or based on tumour volume), mice were enrolled into treatment cohorts and dosed according to the allocated treatment schedules. For trial design 2 in which treatment started at a fixed tumour volume, mice were assigned into cohorts using a stratified randomisation approach which aims to evenly distribute initial tumour volume sizes. To achieve this, tumour volumes from a group of mice are ranked from low to high and then assigned to a cohort using a spiral approach (e.g. 1-3, 3-1, 1-3... for three cohorts).

For CT and CTO trial arms, four cycles of therapy were administered (three weeks per cycle) to total 11 weeks of treatment. Carboplatin was administered by intravenous injection once every three weeks (day 1 of each cycle with 0.16mg and 40mg/kg doses for trial designs 1 and 2 respectively). Paclitaxel was administered by intravenous injection once weekly (days 1, 8, 15 of each cycle with doses of 0.07mg and 7mg/kg for trial designs 1 and 2 respectively). Olaparib was administered by oral gavage on days 3-14 of each cycle (0.05mg or 50mg/kg for trial designs 1 and 2 respectively). For the olaparib monotherapy, olaparib (50mg/kg) was administered by oral gavage with 5 days on/2 days off for 11 weeks. For AZD1775 monotherapy, AZD1775 (120mg/kg) was administered by oral gavage with 5 days on/2 days off for 11 weeks. For olaparib/AZD1775 combination, models 1006 and 1040 were administered with olaparib (50mg/kg) 5 days on/2 days off and AZD1775 120 mg/kg 5 days on/2 days off. Models 1022 and 1141 were administered with olaparib (50mg/kg) daily and AZD1775 (60mg/kg) 3 days on/4 days off.

Paclitaxel (Selleckchem, S1150) was formulated in 1:1 Ethanol:Kolliphor (Sigma, C5135-500G) and diluted prior to use in Vetivex saline. Carboplatin (Selleckchem, S1215) was formulated in sterile water. Olaparib (provided by AstraZeneca) was formulated in 30% Kleptose/10% DMSO in water and AZD1775 (provided by AstraZeneca) was formulated in 0.05% methylcellulose. Mice were carefully monitored for adverse effects and tumours were measured weekly with callipers. Mice were humanely sacrificed at the defined endpoint (either immediately after treatment or at tumour volume 1500 mm^3^). All *in vivo* drug trials were performed at Cancer Research UK Cambridge Institute animal facility.

### Measurement of tumour volume

Callipers were used to measure the volume of the mouse tumours by using the height (h), and width (w) weekly. The tumour volume (mm^3^) was determined by using the equation:

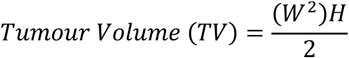

### PDTX tumour growth and drug response modelling

Raw data from trial design 1 was analysed by comparing tumour volumes at defined time points, area under the curve (AUC) and regression coefficient following log2 transformation of tumour volume. AUC was calculated using Prism (v8.1.0), which computes the AUC using the trapezoid rule. In the instance that one cohort reached limits before the end of treatment, the final measured volume was used for the remaining days. The regression coefficient (slope) was calculated by linear regression, following log2 transformation of the tumour volume over time up to the end of treatment. All parameters were compared between cohorts using two-tailed, unpaired Welch’s t-test, not assuming equal variance.

For trial design 2, based on the assumption of exponential tumour growth (35), a random intercept linear mixed model was fitted to each PDTX model to estimate the effect of treatment on the daily growth rate. These models were used to compare growth trajectories between drug-treatment arms and untreated. The linear mixed model was fitted on the log volume of the tumour on day *i* after the start of the treatment, and the effect on the treatment and a random slope for every mouse was included, allowing us to model the variability in growth rates observed on different replicates. P-values were corrected to multiplicity adjustments for within-mouse comparison and analysis. In addition, it was only included the period under treatment in our models.

For each model, the following metrics were derived to quantify the drug treatment effect:

1. The mean difference in daily growth rate between the treatment group and the untreated group, estimated with the interaction between growth rate and treatment group in the linear mixed model.
2. The area under the curve between the estimated marginal mean growth of the treated and the untreated groups.
3. Mean daily growth rates for the treatment groups.
4. The predicted volume for each mouse at the end of treatment was derived from the linear mixed model at days 84.

### DNA/RNA extraction from PDTX tumours for bulk RNA and whole exome sequencing

PDTX tissue samples were flash frozen in liquid nitrogen. Tissue samples were homogenised using a tissue lyser (Qiagen). DNA and RNA for bulk RNA-sequencing and whole exome sequencing (WES) was extracted using AllPrep DNA/RNA kit (Qiagen) following the manufacturer’s instructions.

### Bulk RNA-sequencing

RNA-Seq libraries for Illumina sequencing were prepared using TruSeq stranded mRNA kit (Illumina, 20020595) according to manufacturer’s instructions for high-throughput sample workflow. An input of 500 ng of total RNA per sample was used for library preparation. After DNA fragments were enriched by PCR, all libraries were quantified using KAPA Library Quantification Kit (KAPA Biosystems) and analysed by TapeStation (Agilent). Successful libraries were normalised based on concentration and pooled. Libraries were sequenced on either HiSeq 4000 or NovaSeq 6000 sequencing platforms (Illumina) according to the manufacturer’s instructions. Sequencing was performed using either 50 bp single-end (SE) reads or 50 bp paired-end (PE) reads, to generate on average 15 million total reads per library. The bioinformatics analysis of the RNA-seq data was perform using R 4.0.3. Gene Set Enrichment Analysis (GSEA) was done in single-sample mode (ssGSEA) with the GSVA package and fgsea package for pathway analysis. Normalization and differential gene expression analysis was done using edgeR package.

### Whole exome sequencing (WES) sequencing

WES sequencing libraries were prepared using Nextera Flex for Enrichment (Illumina, 20025524) according to manufacturer’s instructions. Libraries were sequenced on the NovaSeq 6000 using 100 bp paired-end reads, aiming for approximately 100x coverage. Reads were aligned using novoalign (Novocraft) applying our ICRG pipeline to remove mouse contamination (58). Bam files were merged, sorted and indexed using samtools. Duplicates were marked using Picard tools. Candidate SNVs and Indels were called using GATK HaplotypeCaller. Variants were annotated using annovar (version 2018-04-16) for gene/exon annotation, 1000 genomes (version 2015aug), dbSNP (version snp142). Somatic mutations were identified by filtering out calls present in dbsnp, 1000 genomes or in an internal panel of normal (n=95). In the latter, variants were called with the same pipeline and reported if present in at least 2 samples.

### Preparation of single-cell suspensions for mass cytometry and single-cell RNA-sequencing

Cryopreserved PDTX tumour fragments were thawed rapidly into RPMI (Gibco, Invitrogen) and mechanical and enzymatic dissociation was performed using the medium tumour dissociation protocol on a GentleMACS Dissociator, using the human tumour dissociation kit (Miltenyi, 130-095-929) according to manufacturer’s instructions. After tissue dissociation, single-cell suspensions were filtered through 70 μm meshes (BD Biosciences) and transferred into RPMI (Gibco, Invitrogen).

### Single-cell RNA-sequencing and processing

Single cell suspensions of PDTX tumours were prepared as described above. Single-cell sequencing was performed using the Chromium Single Cell 3’ GEM, Library & Gel Bead Kit v2 (10x Genomics, PN-120237) and Chromium Single Cell A Chip Kit (10x Genomics, PN-120236) according to the manufacturer’s instructions. Libraries were sequenced by HiSeq2500 (Illumina) or by NovaSeq 6000 SP or S1 flow cells (Illumina) with paired-end reads and dual-indexing. Cellranger v5.0 (10x Genomics) was used to generate raw count matrices with the hgmm reference. CellBender (59) was used to remove ambient noise and generate the count matrices for the filtered cells. Cells with at least 750 UMIs from non-mitochondrial genes and less than 40% UMIs from mitochondrial genes were kept for analysis. Next, Souporcell (60) was used with K=2 to cluster cells into human and mouse clusters and to remove doublets. Cells in the human cluster with the majority of their UMIs coming from human genes were kept. Finally, Scrublet (61) was used with 0.3 as a cut-off for doublet score to further remove potential doublet cells.

### Single-cell RNA-sequencing Metacell analysis

To select feature genes for MC analysis we first identified high variance and strong genes, then removed blacklisted genes (mitochondrial and a few strong non-coding genes, and gene modules correlated with cell cycle, interferon and stress responses, supplementary table 2) which resulted in a set of 3,390 feature genes. MCs were derived as described (39) using K = 110, standard bootstrapping and MC splitting. We further filtered 2 MCs suspected to be nuclei stripped from their cytoplasm as they had high fraction of UMIs from lncRNA and mitochondrial genes, low number of UMIs and low fraction of UMIs from ribosomal genes. We also filtered out a single MC suspected to contain Epithelial – Mesenchymal doublet cells. The derived final model included 21,087 cells partitioned into 153 MCs.

### Mass cytometry processing

Time-of-flight mass cytometry (CyTOF) was performed following the optimised protocol for breast cancer PDTXs as described in (20). Briefly, single cell suspensions of PDTX tumours were prepared as described above and cells were exposed to the intercalator Rhodium (103Rh), a live-dead exclusion marker for CyTOF (Fluidigm, 201103). Single cell suspensions of PDTXs were individually subjected to fixation (in 2% PFA), permeabilization and palladium barcoding using the Cell-ID™ 20-Plex Pd Barcoding Kit (102Pd, 104Pd, 105Pd, 106Pd, 108Pd, and 110Pd) as per manufacturer’s instructions (Fluidigm, 201060). After barcoding, 20 samples were pooled into one tube; biological replicates were split between pools and one common sample was included in each pool for downstream analysis. Cell suspensions were incubated with a mix of the extracellular antibodies in cell staining buffer (CSB) for 30 min at room temperature, followed by incubation of cells with 100% ice-cold methanol at 4 °C for 15 min. Cells were incubated with intracellular antibody mix for 30 min at room temperature, followed by secondary antibody mix for 20 min at room temperature. Several wash steps were performed between each step using CSB. Cells were then incubated with the intercalator Iridium (191Ir and 193Ir) (Fluidigm, 201192), an intact single-cell inclusion marker, according to manufacturer’s instructions. EQ four-element calibration beads (140/142Ce, 151/153Eu, 165Ho and 175Lu) were added to the sample as per manufacturer’s instructions (Fluidigm, 201078) and samples were run on the HELIOS instrument at a concentration of 0.6–1 million cells per ml.

### Mass cytometry analysis

Raw data were normalised with the EQ Calibration beads and individual sample de-convolution (using the barcodes) was performed with the commercially available software (Fluidigm). Initial data quality and gating for intact single and alive cells were determined using traditional cytometry visualisation as described in (20).

Data analysis was performed using software available from Cytobank (38). Data from all channels were transformed using the hyperbolic arcsinh function. ViSNE plots were created using all markers except ER, CK8/18, HER2 (not expressed), and event counts were down-sampled to the lowest sample. For ViSNE analyses, the following settings were utilised: 1000 iterations, 30 perplexity, 0.5 theta. Human cells were manually gated using a combination of human-reactive antibodies against CD298 and EpCAM, and mouse-reactive antibodies against MHC class I and CD45. Human cells were further gated for the CD298+/EpCAM+ population. FlowSOM clustering was performed on human cells using hierarchical consensus clustering method, 7 metaclusters, 100 clusters, 10 iterations and normalised scales. FlowSOM clustering was performed using only expressed cell-type markers: EpCAM, CD44, VE-Cadherin, CD49F, EGFR, Vimentin.

### Western blot

Flash frozen PDTX tissue samples were homogenised using a tissue lyser (Qiagen) in protein lysis buffer: Tris-HCl (50 mM), sodium chloride (150 mM), TritonX-100 (1%), EDTA (5 mM), sodium fluoride (50 mM), β-Glycerophosphate (25 mM), protease inhibitors (ThermoFisher, 1861279), phosphatase inhibitors (ThermoFisher, 78427). Tissue was incubated on ice for 30 minutes, and protein-containing supernatants were collected by centrifugation at 4°c for 30 min. Bicinchoninic acid assay (BCA protein assay kit, Pierce) was used to quantify proteins. Lysates (20 μg of total protein) were run on 4-12% polyacrylamide gels (NuPAGE, Invitrogen) by SDS-PAGE, transferred to nitrocellulose membranes, blocked in Tris-buffered saline/0.1% Tween (TBST)/5% milk and probed with antibodies against E-Cadherin (Cell Signaling Technology, 3195S, 1:1000 dilution), Vimentin (Cell Signaling Technology, 5741S, 1:1000) and β-actin (Sigma-Aldrich, A5441, 1:5000). All primary antibodies were diluted in TBST/5% milk and incubated overnight at 4°c. Membranes were washed in TBST and probed with secondary antibodies: anti-rabbit HRP (Dako, P0448, 1:2000) for E-Cadherin and Vimentin, and anti-mouse HRP (Dako, P0447, 1:2000) for β-actin. All secondary antibodies were diluted in TBST/5% milk and incubated for 1 hr at room temperature. For BRCA1 western blot, protein was lysed in RIPA buffer, run on 3-8% gel (NuPAGE, Invitrogen) and probed using BRCA1 (Santa Cruz, sc-6954, 1:200) and β-tubulin (Cell Signaling Technology, 2146S, 1:1000) primary antibodies, and anti-mouse HRP (Dako, P0447, 1:2000) and anti-rabbit HRP (Dako, P0448, 1:2000) secondary antibodies. Enhanced chemiluminescence (ECL) was performed by incubating membranes with Tris-HCl (100 mM), luminol (1.25 mM), P. Coumaric acid (0.2 mM), hydrogenic peroxide (0.009%) diluted in distilled water for 1 minute at room temperature. Membranes were developed by exposing x-ray films to emitted light.

### Immunohistochemistry

Immunohistochemistry was performed as described in (19). Briefly, tissue samples from PDTXs were fixed in 10% neutral buffered formalin, embedded in paraffin and subsequently used to extract 0.6 mm cores for TMAs construction. TMA immunohistochemical staining was performed on 3-μm-thick sections that had been de-paraffinised and rehydrated on the automated Leica ST5020 system. Appropriate antigen retrieval treatment was performed, followed by antibody staining using Polymer Refine Detection System (Leica, DS9800) using the automated Bond-III platform. Heat-induced antigen retrieval (sodium citrate or Tris-EDTA) was performed at 100°c. Enzymatic antigen retrieval utilised Bond enzyme concentrate (Leica, AR9551) (101.8μg/mL proteolytic enzyme concentration) at 37°c. The signal was enhanced using DAB Enhancer (Leica Biosystems) for all antibodies apart from ER and PR. After staining, sections were dehydrated, cleared in xylene on the automated Leica ST5020 and mounted with the CV5030 (Leica). HER2 staining was performed separately using the PATHWAY rabbit anti-HER2 monoclonal antibody (Ventana, 790-2991) and the iView DAB Detection Kit (Ventana, 760-091).

### PDTX *ex vivo* drug screening

Cryopreserved PDTX tumours were dissociated into single cell suspensions as described above, and resuspended in cell culture media: RPMI-1640, supplemented with serum-free B27 (100x), EGF (20 ng/mL), FGFβ (20 ng/mL), Penicillin-Streptomycin (50 U/mL), Gentamicin (5 ug/mL). Cells were plated in 384-well plates with 50 uL per well with cell concentrations of 1-3.0 X 10^6^ cells/mL and were cultured 24 hours prior to dosing. Cells were treated with the indicated compounds in triplicate using a 7-dose response curve. All drugs except AZD1056 were dosed at 10, 3, 1, 0.3, 0.1, 0.03, 0.01 uM. AZD1056 was dosed at 1, 0.3, 0.1, 0.03, 0.01, 0.003, 0.001 uM due to solubility. Viability was measured by Cell-Titer-Glo-3D (CTG-3D) (Promega, G9683) 14 days after dosing.

PDTX drug responses were analysed as previously described in (19). Briefly, the observed response was computed as 100 - (100 * (intensity-negative control)/(positive control - negative control)). Non-parametric isotonic regression using the R function isoreg was used to fit the set of technical replicates of a given drug response for a given sample. The area under the curve (AUC) was computed on the model fits using the trapezoid rule with the R package flux.

### PDTX *ex vivo* synergy testing

Cells were prepared and plated as above, and cultured for 24 hours prior to dosing. Cells were dosed in a 7-dose response matrix of the two compounds (10, 3, 1, 0.3, 0.1, 0.03, 0.01 μM) with three technical replicates per dose combination. Dose response was also assessed for each drug as a single-agent. Viability was measured by CTG-3D (Promega, G9683) 7 days after dosing. Synergistic effects were measured using the Bliss model, which compares the observed response under a given combination of two drugs and concentrations, and the predicted response under a model of independence (49). Single-agent responses were used to calculate predicted responses using the following formula, where R_A_ is the response at a given concentration of drug A and R_B_ is the response at a given concentration of drug B. R_predicted_ = R_A_ + R_B_ - R_A_R_B_. This was compared to observed responses (R_observed_), which were fitted using a bivariate isotonic fit on the drug concentration obtained from the R package isotonic.pen (v1.0). The synergistic score was calculated by subtracting R_predicted_ from R_observed_. A combination was considered synergistic if R_observed_ > R_predicted_ and antagonistic if R_observed_ < R_predicted_.

## ACKNOWLEDGEMENTS

This research was supported with funding from Cancer Research UK (Caldas Core Grant A16942 and CRUK Cambridge Institute Core Grant A29580), and an ERC Advanced Grant to C.C. from the European Union’s Horizon 2020 research and innovation programme (ERC-2015-AdG-694620). A.B. is supported and receives funding from the Institute of Cancer Research (ICR), London. O.M.R. is supported by NIHR Cambridge Biomedical Research Centre (BRC-1215-20014) and the Medical Research Council (United Kingdom; MC_UU_00002/16). Y.E.L. is funded by the European Union’s Horizon 2020 research and innovation programme under the Marie Sklodowska-Curie grant agreement No 895808. D.G.R. is funded by the Cambridge Commonwealth, European & International Trust. S.W. was supported through the Fulbright U.S. Student Program, which is sponsored by the U.S. Department of State and the U.S.-U.K. Fulbright Commission. We thank AstraZeneca for providing materials used in this project, and for information about drug formulation and administration. We thank the Cancer Research UK Cambridge Institute Core Facilities (Biological Resources Unit, Preclinical Genome Editing, Genomics, Bioinformatics, Histopathology, Flow Cytometry, and Biorepository) for support during the execution of this project. We would like to acknowledge and thank Ankita Sati who contributed to the development of the PDTX biobank in the Caldas group in Cambridge.

We would like to thank all the patients who participated in the PARTNER clinical trial and who donated their samples for generation of PDTXs. We also thank the Cambridge Cancer Trials Centre PARTNER co-ordination and clinical operations teams, at Cambridge University Hospitals and all the participating sites in the PARTNER UK collaborative. PARTNER was supported by Cancer Research UK and AstraZeneca; J.A. is supported by the CRUK-Cambridge Centre. Grants: CRUKE/14/048 PARTNER (A20735). Clinical Trials Numbers: EuDRACT No: 2015-002811-13; CTA No:24551/0024/001

We would like to thank the Personalised Breast Cancer Programme (PBCP) and the staff of Precision Breast Cancer Institute for their collaboration and support in helping identify and collect appropriate samples and clinical data for this project. Ethics (IRAS project ID): 246732 East of England - Cambridge Central. Funders: CRUK (A27657); Mark Foundation; Addenbrookes Charitable Trust.

This research was supported by the NIHR Cambridge Biomedical Research Centre (BRC-1215-20014*) and the Cambridge Clinical Trials Unit (CCTU). The views expressed are those of the authors and not necessarily those of the NIHR or the Department of Health and Social Care.

## CONTRIBUTIONS

A.B. and C.C. conceived the study. All experiments were done in C.C’s lab, and analysis of the data in A.B., O.M.R and C.C. labs. A.S. and A.B. designed all the *in vivo* and *ex vivo* experimental approaches. A.S. performed all *ex vivo* experiments, generated molecular data, contributed to the analysis and interpretation of the data and the writing of the manuscript, along with A.B. O.M.R and C.C. performed major reviewing and editing of manuscript. A.B., O.M.R., and C.C. contributed to the design of all experiments and data interpretation. Y.E.L. processed and analysed single-cell RNA-sequencing data, and A.M. contributed to this analysis. D.G.R. and O.M.R. developed strategy to, and performed, linear modelling to analyse *in vivo* data. R.M.G. processed and analysed bulk RNA-sequencing data. M.C., A.G. and R.M. processed and analysed WES data, and A.N. contributed to this analysis. O.M.R. oversaw all computational aspects of this project, contributed to the statistical analysis and interpretation. A.S., D.G., W.G., E.E., M.O., G.L. and A.B. contributed to the methodology, investigation, validation, and resources. D.G. designed the MC panel, and D.G., S.W. and A.S. generated MC data. H.B. and E.P. contributed to the generation of TMAs and IHC data. L.Y., S.K., and Y.C. performed *in vivo* experiments and contributed to *in vivo* experimental design together with A.S and A.B. L.G. set up and ran the PARTNER clinical study, and K.McA. consented patients. J.K. collected samples and organised USS and follow up for PBCP, and J.L. curated data for PBCP under supervision of J.A. J.A. is Principal Investigator of the PARTNER clinical trial and contributed to the clinical interpretation of the data, together A.S., A.B. and C.C. All authors revised the manuscript and approved its final version. A.B., O.M.R. and C.C. were responsible for the supervision and project administration.

